# Noise Gated by Dendrosomatic Interactions Increases Information Transmission

**DOI:** 10.1101/163261

**Authors:** Richard Naud, Alexandre Payeur, André Longtin

## Abstract

We study how noise in active dendrites affects information transmission. A mismatch of both noise and refractoriness between a dendritic compartment and a somatic compartment is shown to lead to an input-dependent exchange of leadership, where the dendrite entrains the soma for weak stimuli and the soma entrains the dendrite for strong stimuli. Using this simple mechanism, the noise in the dendritic compartment can boost weak signals without affecting the output of the neuron for strong stimuli. We show that these mechanisms give rise to a noise-induced increase of information transmission by neural populations.

## I. INTRODUCTION

Biological systems process information with high efficiency despite a machinery characterized by high intrinsic variability. This incongruity may be resolved by considering the beneficial effects of noise on information transmission. At multiple time scales, noise can enhance information transmission for either subthreshold [1–3] or suprathreshold signals [4], a phenomenon known as aperiodic stochastic resonance. Yet, it is unclear how and to what extent neuronal populations exploit the noise inherent to the biophysics of membranes.

Intrinsic noise is known to contribute to the activity of single neurons [5–8]. It is thought to arise from stochastic changes in ion channel conformations regulating vesicle release and action potential generation. If such intrinsic noise were to play a constructive role in information transmission either by boosting subthreshold signals or decorrelating individual elements [1, 4, 9], its intensity should be tuned to the particular input [10, 11]: too little noise does not significantly enhance information, too much degrades it. It is unclear if single neurons can tune the intensity of intrinsic noise, and if so with what precision. Preferably, neurons would have a mechanism to gate noise selectively according to the strength of the input. Here we study the extent with which noise in active dendrites affects the information transmitted by the cell body.

Dendrites are characterized by small compartment sizes [12–14], large intrinsic noise [15–17], and refractoriness [18, 19], which limits their maximal firing frequency. Conversely, the cell bodies of neurons are characterized by large compartment sizes, weak intrinsic noise and can sustain high firing frequencies. These two types of neural subunits are active since they may generate spikes locally [13, 14, 18, 20]. They are also coupled: a dendritic action potential can force a spike in the cell body and the back-propagating action potential couples the compartments in the reverse direction [12–14, 18]. In this article, we show how these features can perform noise gating, and how this leads to an enhancement of time-dependent information transmission.

## II. THE DENDRITE-SOMA SYSTEM

### A. Simplified Biophysical Description

The role of dendrites can be addressed by a simplified biophysical model with a single dendritic compartment connected to the cell body. This dendrite-soma system is typically modelled with resistive coupling between the dendrite and the soma and a reduced set of ion channels on both compartments [21–23]. In an instantiation of such a system (see Appendix), we simulated the response to a constant input delivered with equal strength to both compartments. In addition to this constant component, and to take into account noisier dendritic dynamics, the compartments were stimulated with independent noise scaled to have a thirty-fold higher amplitude in the dendrite (see Appendix). Figure 1 shows that the response consists of short and stereotypical action potentials in the soma, which are often associated with a broader action potential in the dendrite, consistent with experimental observations [13, 18, 20] and detailed compartment modelling [20, 24, 25]. When the depolarizing input was weak, we found that a dendritic spike would consistently precede the somatic spike (Fig. 1 (a) and (c)). In contrast, when the depolarizing input was strong, the dendritic spike would generally follow the somatic spike (Fig. 1 (b) and (c)). In addition, we observed that the firing rate of the two-compartment system would follow more closely the firing rate of an isolated dendritic compartment when the input was weak, and more closely the firing rate of an isolated somatic compartment when the input was strong (Fig. 1 (d)).

**FIG. 1.**
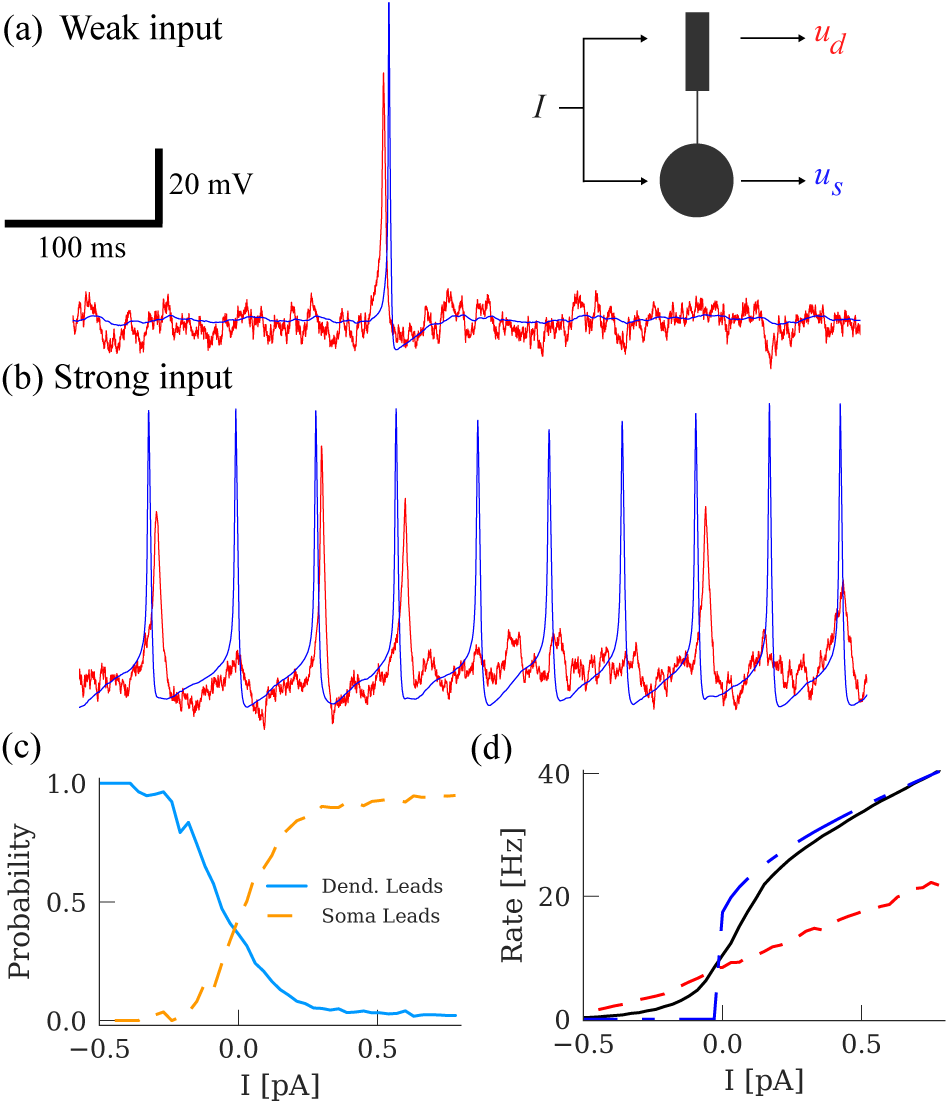
Exchange of leadership in a simplified biophysical model of the dendrite-soma system. (a) The dendrite-soma system (see Appendix) receiving a weak input in both the dendrite and the soma. The membrane potential of the soma (blue trace) and the dendrite (red trace) shows action potentials initiated first in the dendritic compartment. (b) For a strong input, the somatic compartment fires regularly and generally leads the dendritic action potentials. (c) The fraction of somatic spikes that were preceded by a dendritic spike within 8 ms (blue, full line) and the fraction of dendritic spikes that were preceded by a somatic spike by at least 8 ms (orange, dashed line) are shown. Spike timing is taken to be the time of crossing -30 mV from below. (d) The firing rate of the coupled system is shown as a function of the input strength (black, full line). Isolating the compartments by fixing the coupling conductance to zero shows that the dendrite-soma system interpolates between the isolated dendritic compartment (red, dashed line) and the isolated somatic compartment (blue, dash-dot line).

### B. Integrate-and-Fire Description

In order to identify central mechanisms from biophysical models, we used a simple yet accurate abstraction.

We considered that the tip of dendrites can emit stereotypical spikes associated with a relative refractory period longer than that at the soma. We therefore modeled a dendrite-soma system as two interconnected integrateand-fire units, a system studied in the context of connected pairs of neurons [26, 27]. To model dendrite-soma systems, we considered that each compartment has independent intrinsic noise, a distinct refractory period and common stimulation of intensity *s*. These effects are distinct from dendritic NMDA-spikes [25, 28–32], of calcium spikes [21, 22, 33] or other simplified models of dendritic activity lacking either a back-propagating action potentials or a clear refractory period [34, 35].

The dendritic (somatic) potential *u*_*X*_ (*u*_*Y*_) evolves according to the Langevin equations

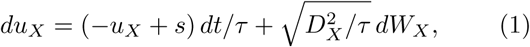

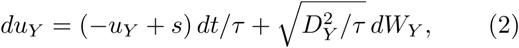

with membrane time constant *τ* [36]. Upon reaching a threshold at *u*_*X*_ (*u*_*Y*_) *>* 1, a unit is said to fire. When a unit fires, it causes a strong potential jump *b* in the other unit, consistent with strong active coupling between soma and dendrites [12–14, 20]. This jump is implemented numerically in the timestep after the firing time. After spiking, a unit remains clamped to the reset potential *V*_*X*_ (*V*_*Y*_) for unit *X* (*Y*) during an absolute refractory period *T*_*R*_, after which the membrane potential follows Eqs.1-2. We model a different relative refractory period with different reset potentials since it takes a longer time to relax from a lower reset. We choose *V*_*X*_ *< V*_*Y*_ to model a longer relative refractory period for the dendrite. Lastly, each unit is subjected to an intrinsic noise denoted by the independent Wiener increments *dW*_*X*_ and *dW*_*Y*_ in Eqs. 1-2 with intensity scaled by *D*_*X*_ (*D*_*Y*_). To comply with the intensity of intrinsic noise expected in neocortical dendrites[15–17], we consider that unit *X* is noisier than unit Y (*D*_*X*_ *> D*_*Y*_) and study the dynamics of the system when *D*_*X*_ is varied within a realistic range. Our analysis does not include an explicit subthreshold coupling between the compartments reflecting weak electrotonic coupling in the presence of active spike propagation in cortical dendrites [12–14]. With these parameter restrictions, the noisier and more refractory unit (X) models an active dendrite while unit Y corresponds to the soma, and so it is the output of the system.

## III. EXCHANGE OF LEADERSHIP AS NOISE GATING

### A. Exchange of Leadership

The coupling between the units implies that whenever one of the units fires, the other has a high probability to discharge immediately afterwards. The relative refractory period prevents another firing event to directly follow this dual firing. Between these dual spiking events the units are effectively independent. Then, the first unit reaching threshold dictates the firing dynamics of the coupled system. In the subthreshold regime, i.e. for *s <* 1, the potential of each unit cannot cross threshold without noise and both *u*_*X*_ and *u*_*Y*_ would saturate to *s* in this case. At this potential, the noisier unit has a greater probability to fire since *D*_*X*_ *> D*_*Y*_. The noisier unit *X* will fire more often and therefore be the *leader*, entraining the more deterministic unit *Y* as illustrated in Fig. 2 (a). On the other hand, in the suprathreshold regime (*s >* 1), both units fire without noise. Hence, following a spike, unit *Y* can take advantage of its higher reset value and cross threshold before *X* (Fig. 2 (b)).

**FIG. 2.**
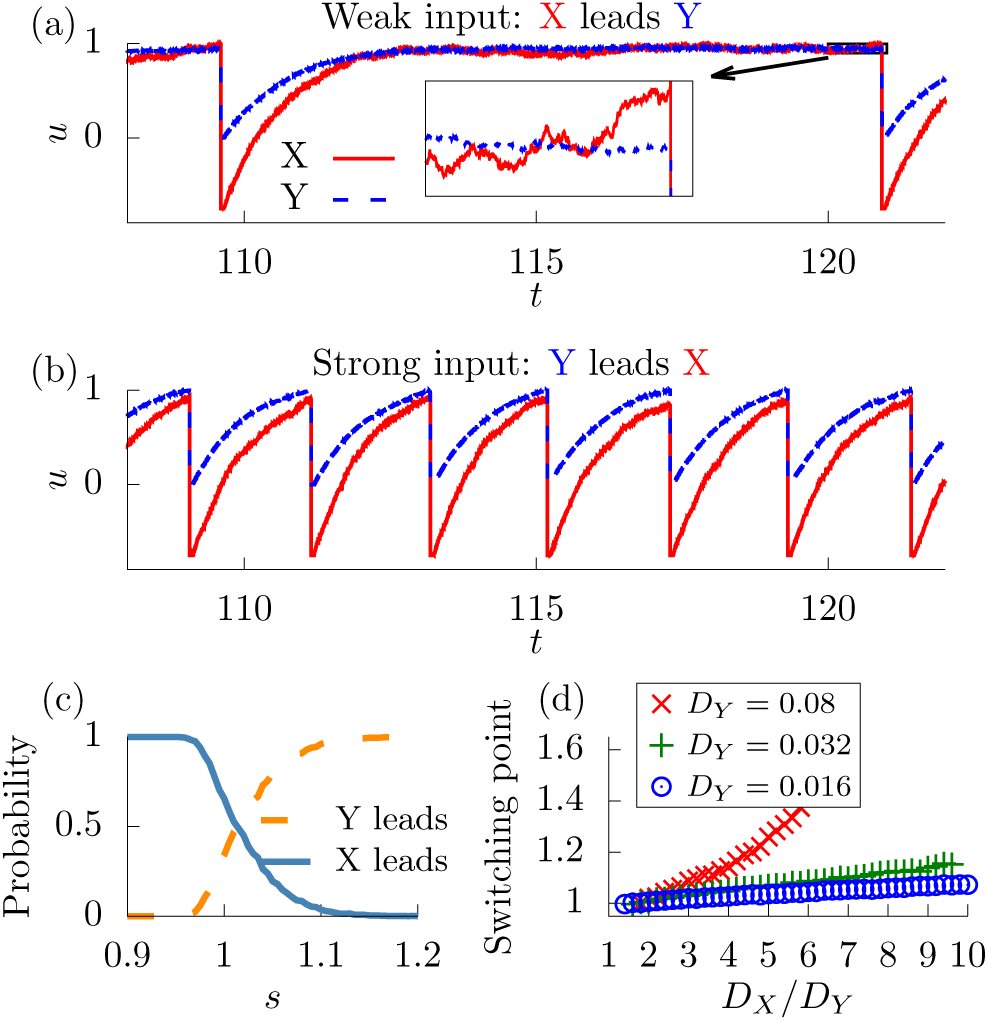
Noise gating in coupled integrate-and-fire units. (a) Membrane potential response of Y (somatic, blue dashed line) and X (dendritic, red full line) units for *s* =0.95 and *D_X_* = 3*D_Y_*. Inset: expanded view near threshold, where the Xunit crosses first (arrow). (b) Same as (a) but for stronger input *s* = 1.15. (c) Entrainment probability for a range of input strengths, as described in the caption to Fig. 1 The time window used to establish leadership in the integrateand-fire model was of the same order as the integration time step. The input intensity where entrainment probabilities cross is the switching point. (d) Dependence of the switching point on *D_X_/D_Y_* for different values of *D_Y_*. Parameters are described in the reference section [36].

An exchange of leadership, or *switching*, occurs as *s* is varied from subthreshold to suprathreshold values. We computed the probability that *X* entrains *Y, P*_*X→Y*_, by counting the fraction of *X* spikes that are immediately followed by a *Y* spike. Figure 2 (c) shows *P*_*X→Y*_ and its complement *P*_*Y* *→X*_ calculated by numerical simulation of Eqs. 1-2 in a typical example of switching. The dendritic leadership is complete and sustained at subthreshold input strengths. As *s* is increased, dendritic leadership is diminished and *P*_*X→Y*_ decreases. Simultaneously, somatic leadership is augmented and *P*_*Y* *→X*_ increases. In the strong input regime the exchange of leadership is complete such that *P*_*X→Y*_ reaches zero and *P*_*Y* *→X*_ one.

We observed complete switching provided that the relative dendritic noise *D*_*X*_ is sufficiently greater than *D*_*Y*_ and sufficiently small compared to the coupling amplitude *b* to ensure strong effective coupling. Within this range, the switching point defined by *P*_*X→Y*_ = *P*_*Y* *→X*_ = 0.5 is not constant but increases with *D*_*X*_ (Fig. 2 (d)).

A mismatch of the level of noise across compartments combined with a mismatch of refractoriness can therefore mediate the input-dependent exchange of leadership.

### B. Noise Gating

Since the switching seen at strong inputs implies that the influence of high intrinsic noise in unit X is removed from the output, we remark that the system effectively reduces, or *gates*, noise as a function of input intensity. Furthermore, gating emerges close to the deterministic threshold, precisely at the point where the role of noise switches from beneficial to detrimental in single-unit encoding. It suggests a particular role for *noise gating*: a more deterministic encoding of supra-threshold inputs and a noise-assisted encoding to resolve subthreshold inputs within the same encoding device and without the use of feedback.

## IV. EFFECTS ON ENCODING PRECISION

### A. Stationary Inputs

To show the role of a dendrosomatic mismatch of noise on encoding quality, we investigated the consequences of noise gating on stationary firing statistics (Fig. 3). For various input strengths *s*, we computed the mean firing rate and the variance of the interspike intervals 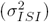. At subthreshold input strengths, the firing rate of the coupled system is identical to the firing rate of an uncoupled X-unit. In this regime, the dendritic compartment controls the timing of the somatic compartment, consistent with in vitro recordings [37] and our biophysical model (Fig. 1 (a)). As the input strength is increased, the firing rate of the coupled system starts to deviate from that of an uncoupled X-unit, reaching the firing rate predicted for an isolated Y-unit when switching is complete (Fig. 3 (a)). Interspike interval variance similarly switches from a variability predicted by the dynamics of an X-unit subthreshold to a variability predicted by the dynamics of an Y-unit suprathreshold (Fig. 3 (b)), as is to be expected from the gating of X-unit noise. Notably, the coupled system follows the strongest firing rate and the smallest variability, concurrently.

**FIG. 3.**
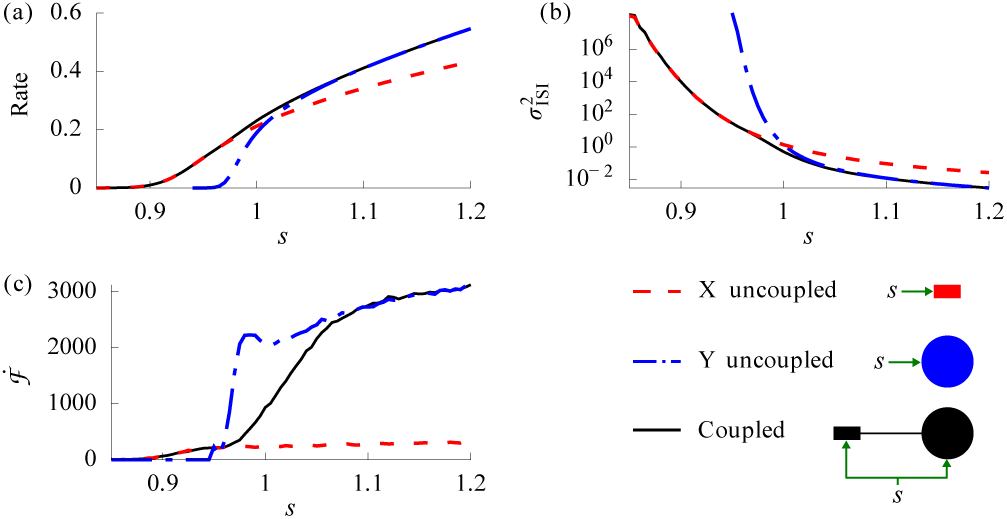
The firing statistics of isolated units constitute asymptotic curves for the dendrite-soma system. (a) Firing rate of coupled system (black line), X-unit alone (*i.e.* dendrite alone, red line) and Y-unit alone (*i.e.* soma alone, blue line) for different input strengths. (b) The interspike interval variance of the coupled system follows the minimum between isolated X and Y units. (c) Fisher information about the input strength. Parameters are described in the reference section [36].

To determine if switching affects signal encoding, we calculated the Fisher information that interspike intervals *T* carry about a constant input. Fisher information measures how sensitive an observable such as the interspike interval *T* is to changes in an input parameter *s*. In practice, we used an approximation to the Fisher information rate

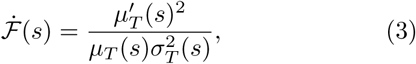

where 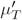 is the derivative of the mean interspike interval with respect to the mean input *s*. The quantity described by Eq. 3 is widely used in studies of neural coding since it is a lower bound on the Fisher information of a population of spiking neurons with independent noise [38, 39].

Consistent with the firing rate and interval variability described above, 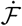 of the coupled system was predicted by X-unit properties subthreshold and by Y-unit properties after complete switching. In the limit of both small and large input strengths, the coupled system showed 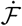 as high as the maximum between isolated X- or Y-units (Fig. 3 (c)). Since for *s* < 1 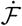 is enhanced by noise, we observe that the stochastic enhancement due to X-unit noise is preserved in the dendrite-soma system. Near threshold, however, the coupled system shows lower 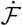 than the isolated Y-unit. To summarize, Fig. 3 suggests that noise gating is beneficial for encoding either weak or strong signals, but not input strengths that lie predominantly close to the deterministic threshold. For these perithreshold inputs, the isolated Y-unit translates small input increments into consistently strong firing rate changes while the coupled system randomly switches between Xand Y-driven firing of with similar rates.

### B. Time-Dependent Inputs

These observations suggest that if a time-dependent input is less often around threshold values, while sampling more consistently both subthreshold and suprathreshold input intensities, then noise gating could enhance the encoding of time-dependent inputs. To achieve this, the coupled system would rely on noisier unit *X* when the input is below the deterministic threshold at *s* = 1. When the input is above the deterministic threshold, the system would switch to a more deterministic encoding by relying on *Y* units. Therefore, we hypothesize that noise gating can enhance encoding of a time-dependent input, even for inputs distributed predominantly above threshold. Additionally, the enhancement should be robust for large dendritic noise since strong noise does not degrade the suprathreshold part of the signal due to noise gating.

To test this hypothesis, we simulated 8000 dendrite-soma systems receiving the same time-dependent input *s*(*t*) with mean 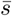 > 1 in the regime of suprathreshold stochastic resonance [4]. An estimate of the population activity is constructed by summing the 8000 spike trains from all Y-units. Encoding quality was quantified by Shannon’s information for the classic channel with additive Gaussian noise [40–42]

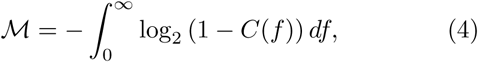

where *C*(*f*) is the coherence between the population activity and the input *s*(*t*) for each frequency *f* [43]. *M* encapsulates the frequency-resolved measure of correlation between input and output fluctuations *C*(*f*) into a single quantity. This quantity is distinct from the average firing rate, and it is used as a lower bound on the mutual information between the time-dependent input and the set of spike trains.

Figure 4 shows *M* for increasing intrinsic dendritic noise and two types of input currents *s*(*t*). For the first type, we considered a Jump-Diffusion Process (JDP) producing random fluctuations around two states with random switching times between the high and the low states. The JDP parameters [44] were chosen to produce a bimodal distribution centered slightly above threshold such that the switching point (Fig. 2 (d)) may cross the center of the input distribution as *D*_*X*_ is varied (Fig. 4 (a)). Although the exact mutual information has not yet been derived for JDPs, this process was chosen to ensure a bimodal distribution of inputs without imposing a periodic structure. It is a physiologically realistic input distribution since sinusoidal sensory inputs and up and down states are frequently treated in the context of neuroscience [45–47]. We compared JDP encoding with encoding of a Gaussian Process (GP) simulated with the Euler-Maruyama method with matched mean and variance (Fig. 4 (b)), focusing, as a first step, on narrow input distributions.

**FIG. 4.**
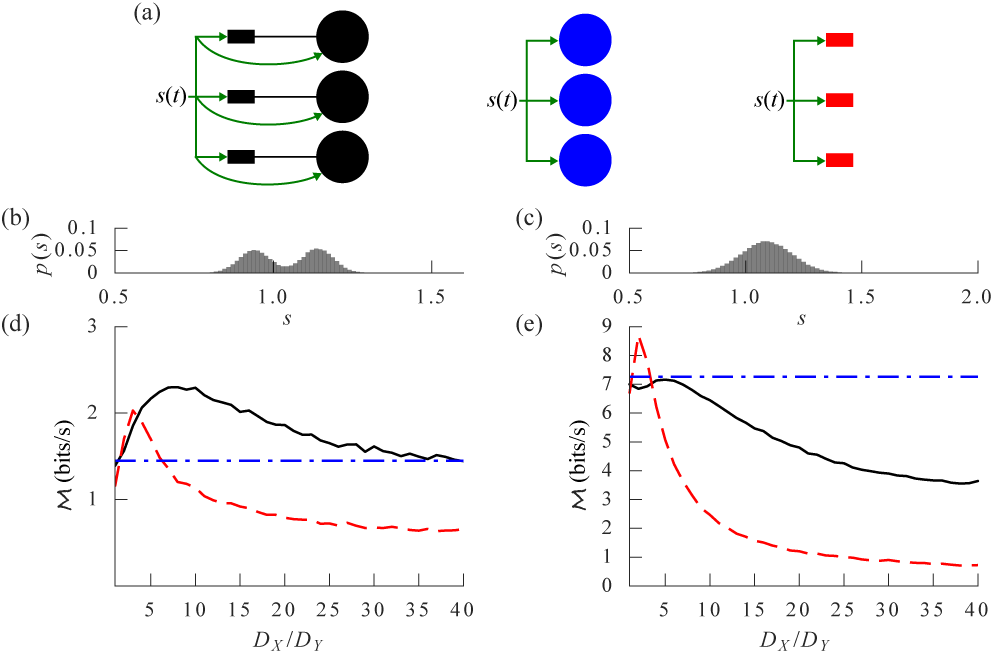
Stochastic resonance for bimodal input distributions. (a) Schema illustrating an neural ensemble made of dendritesoma systems (black), somatic units alone (blue) or dendritic unit alone (red). Input distribution for (b) the JDP and (c) a GP with mean matched to that of the JDP. (d) For the JDP, *M* calculated from the summed activity of 8000 coupled units (black line) surpasses that of isolated Y-units compartments (blue dot-dash line) and of isolated dendritic compartments (red dashed line) for a broad range of dendritic noise. (e) The resonance for a GP is reduced with respect to the resonance of obtained using the JDP (d). We used *τ* = 10 ms to represent *M* in bits per second, see the reference section [36] for all other parameters.

For narrow input distributions, a resonance as a function of *D*_*X*_*/D*_*Y*_ is seen for the JDP but not the GP (Fig. 4 (d)-(e)). Consistent with a more frequent sampling of elevated 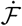 peri-threshold [48], *M* is generally higher for the unimodal input than for the bimodal input. At the maximum, or *resonance*, and only for the bimodal input, the coupled system surpasses in encoding quality a more deterministic population made of isolated Y-units (dashdot line in Fig. 4 (d)-(e)). This *stochastic enhancement* is seen for a large range of intrinsic noise. The coupled system also surpasses a population of isolated X-units with the corresponding intrinsic noise intensity, again only for bimodal inputs. The resonance in Fig. 4 (d) can be un-derstood by recalling that two competing processes result from an increase in *D*_*X*_. On the one hand weaker signals can trigger spikes, which increases the coding range and therefore *M*. On the other hand, the switching point increases, decreasing the range of inputs encoded by the Yunit (Fig. 2 (d)), which decreases reliability and therefore *M*. The output unit (unit Y) representation of bimodal inputs can therefore be enhanced by noise in an auxiliary unit (unit X). The optimal noise results from a trade-off between increasing the coding range and decreasing the coding reliability.

The particular scenario illustrated in *M* Figure 4 (c) corresponds to a coding enhancement with respect to the isolated Y-units of up to 61% in (Fig. 4 (c)). We verified that finite-size effects deteriorate this enhancement by halving the population size. This resulted in a reduction of the enhancement to 42% in *M*. We found that the enhancement in *M* was not detectable for a population of 250 coupled systems, but was present for a population of 500 coupled systems. We verified that the enhancement was maintained by replacing the JDP with a sinusoidal input with matched bimodal peaks. We then verified that the enhancement necessitates a paucity of peri-threshold input strengths by simulating a JDP with a mean increased from 1.04 to 1.14, such that the leftmost peak is close to the deterministic threshold. This manipulation removed the stochastic enhancement. We conclude that intrinsic dendritic noise, when gated by input intensity, may improve *M* for inputs rarely lying close to threshold.

Also, this enhancement is more pronounced in range and amplitude than with classical mechanisms for stochastic resonance [1–4] (see red curve in Fig. 4 (d)).

In the context of neuroscience, the inputs are likely to be broadly distributed. We thus considered a broad GP input distribution with twice the standard deviation of the distribution shown in Fig. 4. We find that even when the average input is above threshold (Fig. 5 (a)), the coupled system shows stochastic enhancement over a large range of intrinsic noise levels (Fig. 5 (b)). Since *M* remains high even at the largest levels of noise we have considered, the dendrite-soma system can tolerate noise coming in addition to noise expected from intrinsic sources [15–17]. Background synaptic noise and finite-size noise are examples of extrinsic noise sources which could contribute to this enhancement.

**FIG. 5.**
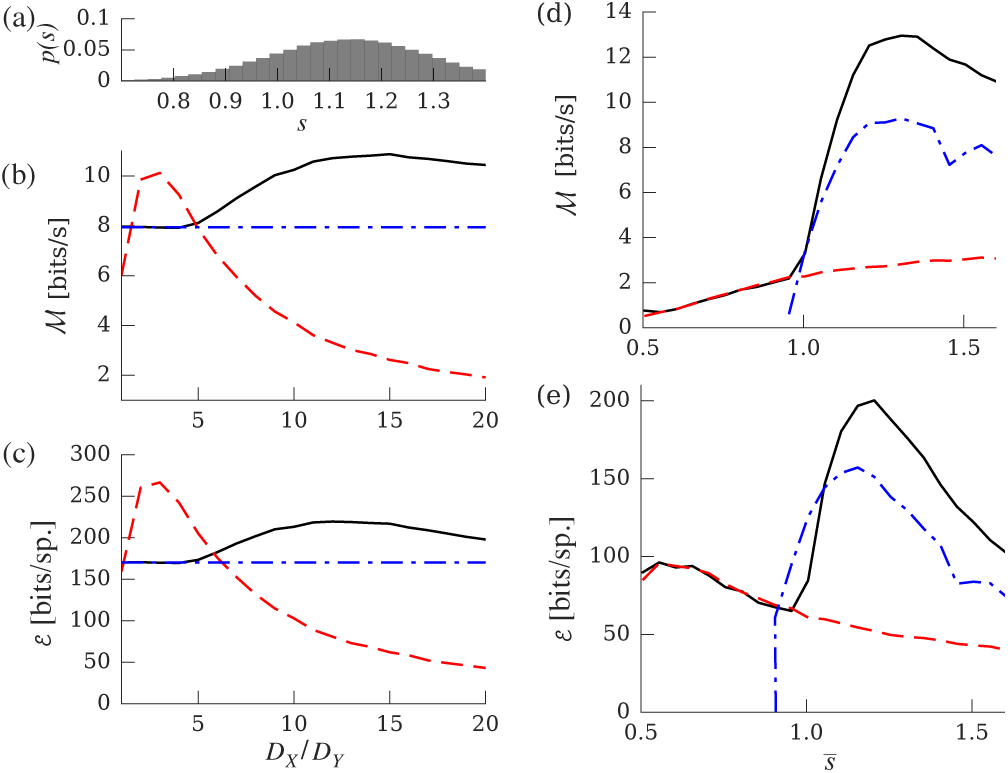
Stochastic resonance for wide input distributions. (a) Input distribution of the GP chosen to cover a large range around a high mean 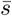. Using the same color code as in Fig. 3, panel (b) shows *M* shows and panel (c) shows *ε* for different *D_X_* calculated with input distribution in (a). We restricted the range for intrinsic noise such that the standard deviation of *u_X_* remains smaller than 0.5, which would correspond to substantial membrane potential fluctuations in physical units. (d) *M* and (e) *ε* for different values of the mean input and for a fixed level of intrinsic noise *D_X_* = 15*D_Y_*. Parameters are described in the reference section [36].

To verify that the information enhancement was not simply due to a rate increase, we normalized *M* by the average firing rate *ν* and considered the quantity

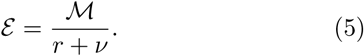

Since every spike comes with a metabolic energy cost, *ε* is interpreted as a measure of energy efficiency of information transfer [49, 50]. We report here a quantity in bits per spike, but this can be converted into bits per ATP molecule using numerical estimates of the number of ATP molecules used for the generation of an action potential [49]. The parameter *r* is interpreted as the firing rate at which the energy expenditure of action potential generation equals the energy expenditure of maintaining a depolarized membrane potential. This parameter is though to vary substantially across neuron types [51], but to be relatively low (we fixed *r* = 5 Hz). Figure 5 (c) shows an enhanced efficiency for large dendritic noise in the coupled system. In addition, the efficiency of the coupled system surpasses the efficiency of either a population of somatic compartments, or a population of dendritic compartments with matched level of intrinsic noise.

We now ask how enhancement depends on the mean input. The *M* of the coupled system matches that of isolated X-units when the mean is subthreshold (Fig. 5 (d)), but surpasses those of isolated X and Y units when the mean is suprathreshold. Therefore, for broadly distributed inputs, noise gating allows stochastic enhancement on a broad range of intrinsic noise levels, thus alleviating the need for precise noise tuning in neurons. The energy efficiency is similarly enhanced (Fig. 5 (e)), but shows two peaks: A first peak subthreshold, matching the enhancement obtained by isolated dendritic units, and a second, much higher peak suprathreshold exceeding the efficiency of isolated somatic units. These results are consistent with the enhanced *M* (Fig. 5 (d)) and a firing rate that follows the largest of either units (Fig. 2). Hence, noise gating in dendrite-soma systems allows an efficient encoding through a stochastic enhancement of subthreshold signals that preserves deterministic encoding suprathreshold.

## V. DISCUSSION

Several other theoretical studies pointed to a functional relevance of spiking or nonlinear summation in dendrites. One view holds that dendrites function as sigmoidal units [28, 29]. This simple description was shown to capture time-average firing rate of biophysical neuron models with detailed morphology and active dendritic conductances supporting NMDA-spikes [25, 31, 52]. Modelling studies have shown that this architecture provides multiple advantages, namely specific sensory computations [53], enhanced memory capacity [30, 35], enhanced dynamic range [54, 55] and flexible gating of specific pathways [32]. These computational advantages are based on a phenomenological description of the time-averaged firing rate, which could remain consistent with the timing-dependent mechanisms described here. Additionally, the timing-dependent network synchrony mechanism discussed in Ref. [34] is likely to hold in the presence of noise gating. Therefore, we add to the known computational advantages of nonlinear dendrites an improvement of encoding precision based on spike-timing interactions.

Experimental studies point to a surprising diversity of active dendrites. The mechanism described here is likely to remain relevant when a combination of long refractory period, small subthreshold coupling and large noise is achieved. We argue that high intrinsic noise and small subthreshold coupling is expected in thin basal dendrites more than one electrotonic constant away from the cell body. First, from measurements of dendrite diameter at the tip of dendrites (0.5-0.6 *μ*m [56, 57]) and experimental estimates of the effective length of a compartment (250 *μ*m [20]), the effective dendritic surface area is estimated to be approximately 400-500 *μ*m^2^. Second, theoretical studies [16] predict that a compartment with surface area of 500 *μ*m^2^ would exhibit noisy fluctuations with standard deviation of 1 mV from the stochastic opening of ion channels, a 10-fold increase from observed somatic noise [58] (consistent with our *D*_*X*_*/D*_*Y*_ ~ 10). Such basal dendrites have accrued refractoriness [18, 19], and longer action potentials [20, 59]. Therefore the known properties of thin basal or oblique dendrites are consistent with the mechanisms described here.

Other dendritic compartments may implement different functions. Notably, the apical tuft of cortical pyramidal cells are known to generate calcium regenerative events, which lead to a burst of action potentials [60, 61] and a dendritic control on somatic gain [62, 63]. In addition, interactions between synaptic plasticity and dendritic dynamics have been the focus of much recent attention [32, 64–66]. In summary, although these mechanisms are not immediately compatible with those described here, they can be combined in a juxtaposition of distinct, spatially segregated compartments within the same dendritic tree.

If noise gating is taking place in the nervous system according to the mechanism described in this article, we predict that spikes encoding a weak stimulus would have been initiated more frequently by the dendrites whereas spikes encoding a strong stimulus would have been initiated more frequently at the soma. Given experimental evidence indicating a connection between the shape of the action potential and the site of initiation [14, 67], it appears possible to test this prediction experimentally. Furthermore, focal pharmacological manipulations can be used to determine the role active dendritic conductances for different strength of sensory stimulation [68, 69].

As a second prediction, we note that the dendritic refractory period should be substantially greater than in the soma for the switching point to be in the typical range of firing rate. Although there are clear indications that the dendrite is more refractory to spiking [18, 19], we are not aware of a direct measurement showing a longer relative or absolute refractory period in the dendrites. A potent discrepancy is likely, given the different composition of ion channels in dendrites [70]. Noting that lengthening of the relative refractory period may arise either from a spike-triggered hyperpolarization or a spike-triggered increase in the action potential threshold [71], an electrophysiology experiment [13, 20, 37, 60] can be designed to measure the difference between somatic and dendritic refractory processes.

## VI. CONCLUSION

The mechanism outlined in this article may provide a functional role of intrinsic noise in active dendrites, where the dendrite act as a noise-assisted encoder, acting only at small firing rates. Also, we have described how the effects of intrinsic noise in a subsystem can be gated by a mismatch of refractory periods between compartments. To conclude, we have outlined a novel mechanism for information enhancement by intrinsic noise. This coding strategy allows to communicate substantially more information per action potential, an interesting approach given the metabolic costs of action potential transmission. The mechanism being simple and easy to implement, it can be inspire novel engineering approaches to signal detection.

## ACKNOWLEDGMENTS

We thank Tilo Schwalger, Jean-Claude Béïque and Len Maler for helpful discussions as well as FRQNT postdoctoral scholarships (R.N.) and the NSERC Discovery Grants (A.L. and R.N.) for funding.

## Appendix Simplified Biophysical Model of the Dendrite-Soma System

We modeled the soma-dendrite system as two connected electrical compartments with different densities of voltage-gated ion channels. We note that this biophysical description is lacking multiple biophysical details, namely multiple dendrites, impedance mismatch between soma and dendrite, multiple other types of ion channels and active or quasi-active propagation. Yet this two-compartment abstraction has been instrumental for understanding many features of dendritic computation [21, 22, 62, 72, 73].

The somatic membrane potential *u*_*s*_ and the dendritic membrane potential *u*_*d*_ evolve according to Kirchoff’s circuit law for the conservation of current

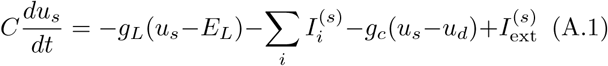

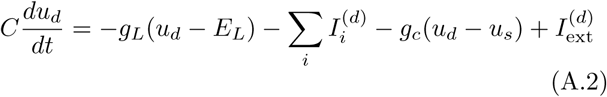

where *C* = 0.75 pF is the compartment capacitance, *g*_*L*_ = 0.2 *nS* is the leak conductance and *E*_*L*_ = *-*70 *mV* is the leak reversal potential. The two compartments are coupled via the coupling conductance *g*_*c*_ = 0.02 nS. Each compartment contains a different set of voltagegated ion channels. The soma is modeled with a combination of inactivating sodium conductance *I*_Na_ and fast rectifying potassium conductance *I*_K_

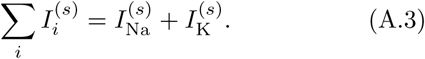

The sodium conductance follows the Hodgkin-Huxley kinetics with an activation gate *m* and an inactivation gate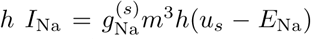, where 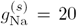 is the maximal conductance and *E*_Na_ = 60 mV is the reversal potential for sodium. The potassium conductance is modeled with a single activation gate *I*_K_ = *g*_K_*n*(*u*_*s*_ − *E*_K_), where *g*_K_ = 10 nS is the maximal conductance and *E*_K_ =*-*70 mV is the reversal potential for potassium. The kinetics of the gating variables *x ϵ {m, h, n}* follows *τ*_*x*_(*u*_*s*_)*ẋ* = *x*_0_(*u*_*s*_) – *x*, with *τ*_*x*_(*u*) and *x*_0_(*u*) described in Table I.

**Table I.**
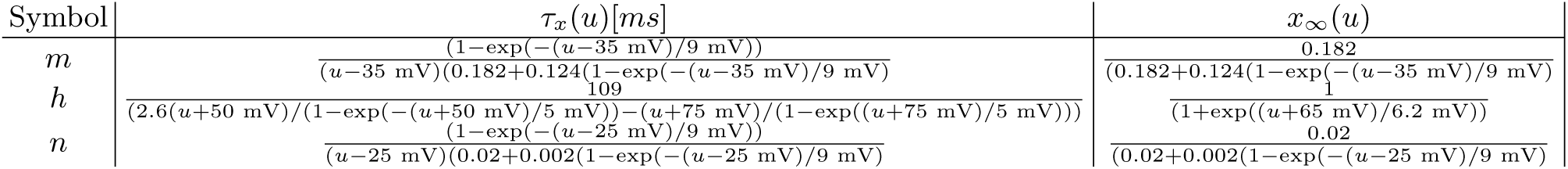
Hodgkin-Huxley kinetics for three types of voltage-gated ion channels. Parametrization follows experiments in neocortical neurons [74–76].

The dendrite contains contains these two types of ion channels, but with lower densities

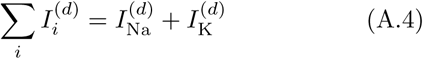

In the dendritic compartment, the maximal conductance of sodium was 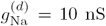 and the that of potassium to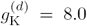 nS. The kinetics of the gating variables *x* ∈ {*m,h,n*} follows *τ*_*x*_(*u*_*d*_)*ẋ* = *x*_0_(*u*_*d*_) – *x*, with *τ*_*x*_(*u*) and *x*_0_(*u*) described in Table I.

Each compartment receives an external input which is partitioned into three terms:

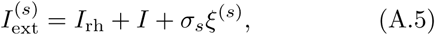

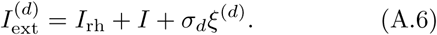

The term *I*_rh_ = 0.57 pA is a constant current corresponding to the rheobase of the system. The additional current *I* controls the current injected into the compartments with respect to this rheobase. We also included independent background noises *ξ*^(*s*)^ and *ξ*^(*d*)^ drawn from a normal distribution (∼ *N* (0, 1)) independently at every time step of size *dt* = 0.1 ms and for each compartment. The noise amplitude was scaled by *σ*_*s*_ = 0.1 pA in the soma, and *σ*_*d*_ = 3 pA in the dendrite.

